# tfboot: Bootstrapping and statistical analysis for transcription factor binding site-disrupting variants in gene sets

**DOI:** 10.1101/2023.07.14.549004

**Authors:** Stephen D. Turner, Kathleen Morrill, Gregory Gedman, Alexander J. Titus

## Abstract

**Motivation:** Genetic variants in noncoding regions can drive changes in phenotype disrupting transcription factor binding site (TFBS) motifs. Other tools including motifbreakR have been developed to assess the impact of TFBS-disrupting variants. Here we introduce the tfboot package for statistically evaluating the TFBS disruption across a *set* of variants in upstream promoter regions.

**Results:** The tfboot package builds on motifbreakR, plyranges, and GenomicRanges to provide methods for bootstrapping TFBS disruption to statistically quantify the impact across gene sets of interest compared to an empirical null distribution. We demonstrate the analysis here on variants in the elephant genome. The tfboot package readily integrates with Bioconductor and tidyverse-based workflows.

**Availability:** The tfboot package is implemented as an R package and is released under the MIT license at https://github.com/colossal-compsci/tfboot.

## 1 Introduction

Transcription factor binding sites (TFBS) are short DNA sequence motifs in the promoter or enhancer regions of genes where transcription factors bind to regulate gene expression. Phenotypic changes can be driven by genetic variants such as single nucleotide polymorphisms (SNPs) which disrupt TFBS (Reshef *et al*., 2018) – indeed the vast majority of human disease-related SNPs found through genome-wide association studies are in noncoding regions (Edwards *et al*., 2013; Buniello *et al*., 2018).

The motifbreakR R package provides methods to ascertain the impact of SNPs disrupting TFBS (Coetzee *et al*., 2015) which function on any genome curated within Bioconductor. motifbreakR evaluates whether the sequence surrounding a SNP is a good match for a TFBS motif drawn from various sources, and assesses how the polymorphism impacts the TFBS motif compared to the wild type sequence.

A common analysis task is evaluating the collective impact SNPs in a *set* of genes or regions. For example, in our research we have a collection of genes known to contribute to body size in various mammalian species. Given a set of genome-wide SNPs from sequencing and a collection of genes of interest, a common question arises: do SNPs upstream of *this gene set of interest* disrupt TFBS more than SNPs in a *randomly selected set of genes*?

Several existing tools provide methods to assess the enrichment of genomic intervals for a given feature, including LOLA (Nagraj *et al*., 2018), and nullranges (Davis *et al*., 2023). Here we introduce the tfboot package, which builds upon the motifbreakR R package to facilitate statistical analysis of SNPs disrupting transcription factor binding sites (TFBS) in *gene sets* using bootstrap resampling to create empirical null distributions.

## 2 Methods

### 2.1 TFBS analysis for SNPs in promoter regions

The tfboot package works with common Bioconductor objects such as GRanges for easy integration with existing Bioconductor-based workflows (Gentleman *et al*., 2004; Lawrence *et al*., 2013), and bootstrapping analysis takes advantage of list-columns in tibbles as implemented in the tidyverse suite of packages (Wickham *et al*., 2019).

A motifbreakR + tfboot analysis starts with a VCF read in with tfboot’s `read_vcf()` function, which is a wrapper around motifbreakR functions for reading VCF files and returns a GRanges object. The `get_upstream_snps()` internally uses plyranges (Lee *et al*., 2019) to take in a list of SNPs as a GRanges object together with a TxDb object (Carlson *et al*., 2016), and returns a GRanges object containing SNPs in the upstream promoter region of genes in the TxDb object.

A standard motifbreakR analysis is then performed on the SNPs in the upstream region of these *k* genes of interest, followed by a motifbreakR analysis on the universe of *all* annotated genes. This step is time-consuming, but precomputing the motifbreakR results on all genes allows for extremely fast bootstrap resampling of *b* bootstrap resamples of *k* genes from this background set. Because all downstream analysis uses standard tidyverse tibbles instead of Bioconductor GRanges objects, tfboot package provides convenience functions to create compact tibbles from the motifbreakR results to reduce disk space and facilitate downstream analysis.

### 2.2 Statistical analysis with bootstrapping

The tfboot package provides downstream functions for summarization, bootstrapping, and statistical analysis of TFBS disruption in gene sets of interest. The tfboot function `mb_summarize` will summarize the results from a motifbreakR analysis into a single-row table with the following columns:

1. `ngenes`: The number of genes in the SNP set.
2. `nsnps`: The total number of SNPs disrupting TFBS.
3. `nstrong`: The number of SNPs with a “strong” effect.
4. `alleleDiffAbsMean` The mean of the absolute values of the `alleleDiff` scores.
5. `alleleDiffAbsSum` The sum of the absolute values of the `alleleDiff` scores.
6. `alleleEffectSizeAbsMean` The mean of the absolute values of the `alleleEffectSize` scores.
7. `alleleEffectSizeAbsSum` The sum of the absolute values of the `alleleEffectSize` scores.

The `mb_bootstrap()` function takes in precalculated motifbreakR results from the universe of all background genes and the number *k* of genes in a gene set to sample, and resamples *k* genes from the precomputed motifbreakR analysis *b* times to create an empirical null distribution of the values calculated above. Finally, the `mb_bootstats()` function takes as input both the summary on the gene set of interest with the bootstrapping results on the background set of genes and calculates p-values comparing the critical value from the user’s gene set of interest against the empirical null. The `plot_bootstats()` function takes the results from this analysis as input to create a visual representation of the critical values against the background null distribution.

## 3 Results

Here we demonstrate a motifbreakR and tfboot analysis on SNPs in Asian elephant (*Elephas maximus*) compared to an African savanna elephant reference genome (*Loxodonta africana*). As *Elephas maximus* morphologically differs from *Loxodonta africana* in ear size and shape, we selected the set of nine genes annotated for outer ear morphogenesis (GO:0042473), as shown in Table 1.

**Table 1.** Nine genes involved in outer ear morphogenesis (GO:0042473) used in this analysis.

| Gene | Name |
| --- | --- |
| PRKRA | Protein kinase, interferon-inducible double stranded RNA dependent activator |
| ZIC3 | Zic family member 3 |
| EYA1 | eyes absent homolog |
| FGFR1 | Fibroblast growth factor receptor 1 |
| GAS1 | growth arrest-specific 1 |
| HOXA1 | Homeobox A1 |
| MAPK1 | Mitogen-activated protein kinase 1 |
| MAPK3 | Mitogen-activated protein kinase 3 |
| TBX1 | T-box transcription factor |

Briefly, we used minimap2 to map PacBio HiFi reads sequenced in the Asian elephant using the African elephant chromosome-level reference genome (Rhie *et al*., 2021) we released earlier this year (NCBI genome accession GCA_030014295.1), and used GATK HaplotypeCaller to call variants run on the Form Bio platform (https://formbio.com/). Asian elephant sequencing data is available on the GenomeArk (https://registry.opendata.aws/genomeark/).

After precalculating the motifbreakR results for upstream promoter regions in all genes in this background set, the tfboot analysis 1,000 bootstrap resamples took approximately 20 seconds on a single CPU on an Apple M2 Macbook Pro. The primary results from the tfboot analysis are shown in Table 2.

**Table 2.** Results from the tfboot analysis with 1,000 bootstrap resamples.

| metric | stat | bootdist | bootmax | p |
| --- | --- | --- | --- | --- |
| nsnps | 2517.000 | (distribution) | 11624.000 | 0.750 |
| nstrong | 1882.000 | (distribution) | 8615.000 | 0.757 |
| alleleDiffAbsMean | 0.811 | (distribution) | 0.834 | 0.468 |
| alleleDiffAbsSum | 2040.547 | (distribution) | 9347.870 | 0.754 |
| alleleEffectSizeAbsMean | 0.181 | (distribution) | 0.188 | 0.622 |
| alleleEffectSizeAbsSum | 454.685 | (distribution) | 2107.027 | 0.754 |

While there may be individual SNPs that disrupt TFBS in promoters of individual genes, the results in Table 2 indicate that there is no statistically significant TFBS disruption in SNPs in the promoter regions of *this set* of genes of interest compared to a null background set of genes of the same size. After calculating bootstrap statistics, the `plot_bootstats()` function can be used to visually display the critical values for motifbreakR TFBS metrics of the gene set of interest against the empirical null distribution (gray), as shown in Figure 1.

**Fig. 1.**
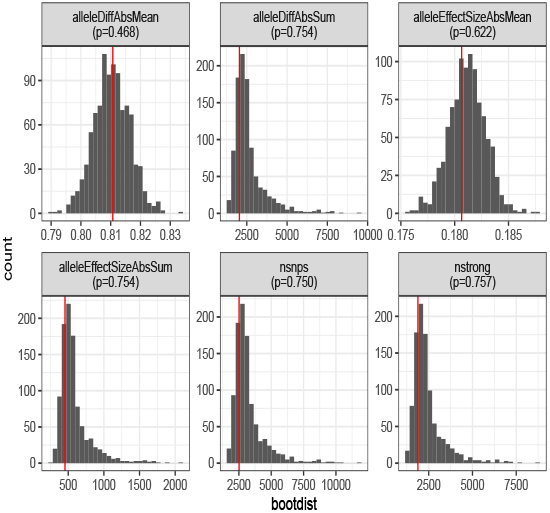
Results from the tfboot analysis with 1,000 bootstrap resamples examining TFBS distruption in the set of nine genes involved in outer ear morphogenesis using SNPs in the Asian elephant relative to the African elephant.

## 4 Conclusion

Here we introduce the tfboot package for statistical analysis of TFBS-disrupting SNPs in gene sets using motifbreakR. We demonstrate an analysis on SNPs between the Asian and African elephant in a set of nine developmentally important genes involved in outer ear morphogenesis, showing that bootstrap resampling and statistical analysis on precomputed motifbreakR results requires <1 minute of compute time. The tfboot package can be run on any genome with available Bioconductor BSGenome and TxDb objects, either publicly available or custom-created from FASTA+GTF files. The tfboot package works with GRanges and other Bioconductor data structures for integration into existing workflows, and results are returned as tibbles with bootstrapping results as nested list columns, facilitating downstream analysis with tidyverse tools. The tfboot package is implemented as an R package and is freely available under the MIT license at https://github.com/colossal-compsci/tfboot.

## Acknowledgements

The authors thank Vijay Kandali and Amanda Kowalczyk for testing and feedback on tfboot package. The authors also thank Brandi Cantarel and Ketaki Bhide for assistance with alignment and variant calling Asian elephant reads against the African elephant reference genome.

## References

Buniello, A., MacArthur, J., Cerezo, M., Harris, L. W., Hayhurst, J., Malangone, C., McMahon, A., Morales, J., Mountjoy, E., Sollis, E., Suveges, D., Vrousgou, O., Whetzel, P. L., Amode, R., Guillen, J. A., Riat, H. S., Trevanion, S. J., Hall, P., Junkins, H., Flicek, P., Burdett, T., Hindorff, L. A., Cunningham, F., and Parkinson, H. (2018). The nhgri-ebi gwas catalog of published genome-wide association studies, targeted arrays and summary statistics 2019. Nucleic Acids Research, 47(D1), D1005–D1012.

Carlson, M. R. J., Pagès, H., Arora, S., Obenchain, V., and Morgan, M. (2016). Genomic Annotation Resources in R/Bioconductor, pages 67–90. Springer New York.

Coetzee, S. G., Coetzee, G. A., and Hazelett, D. J. (2015). motifbreakR: an r/bioconductor package for predicting variant effects at transcription factor binding sites. Bioinformatics, 31(23), 3847–3849.

Davis, E. S., Mu, W., Lee, S., Dozmorov, M. G., Love, M. I., and Phanstiel, D. H. (2023). matchranges: generating null hypothesis genomic ranges via covariate-matched sampling. Bioinformatics, 39(5).

Edwards, S., Beesley, J., French, J., and Dunning, A. (2013). Beyond gwass: Illuminating the dark road from association to function. The American Journal of Human Genetics, 93(5), 779–797.

Gentleman, R. C., Carey, V. J., Bates, D. M., Bolstad, B., Dettling, M., Dudoit, S., Ellis, B., Gautier, L., Ge, Y., Gentry, J., Hornik, K., Hothorn, T., Huber, W., Iacus, S., Irizarry, R., Leisch, F., Li, C., Maechler, M., Rossini, A. J., Sawitzki, G., Smith, C., Smyth, G., Tierney, L., Yang, J. Y., and Zhang, J. (2004). Genome Biology, 5(10), R80.

Lawrence, M., Huber, W., Pagès, H., Aboyoun, P., Carlson, M., Gentleman, R., Morgan, M. T., and Carey, V. J. (2013). Software for computing and annotating genomic ranges. PLoS Computational Biology, 9(8), e1003118.

Lee, S., Cook, D., and Lawrence, M. (2019). plyranges: a grammar of genomic data transformation. Genome Biology, 20(1).

Nagraj, V. P., Magee, N. E., and Sheffield, N. C. (2018). Lolaweb: a containerized web server for interactive genomic locus overlap enrichment analysis. Nucleic Acids Research, 46(W1), W194–W199.

Reshef, Y. A., Finucane, H. K., Kelley, D. R., Gusev, A., Kotliar, D., Ulirsch, J. C., Hormozdiari, F., Nasser, J., O’Connor, L., van de Geijn, B., Loh, P.-R., Grossman, S. R., Bhatia, G., Gazal, S., Palamara, P. F., Pinello, L., Patterson, N., Adams, R. P., and Price, A. L. (2018). Detecting genome-wide directional effects of transcription factor binding on polygenic disease risk. Nature Genetics, 50(10), 1483–1493.

Rhie, A., McCarthy, S. A., Fedrigo, O., Damas, J., Formenti, G., Koren, S., Uliano-Silva, M., Chow, W., Fungtammasan, A., Kim, J., Lee, C., Ko, B. J., Chaisson, M., Gedman, G. L., Cantin, L. J., Thibaud-Nissen, F., Haggerty, L., Bista, I., Smith, M., Haase, B., Mountcastle, J., Winkler, S., Paez, S., Howard, J., Vernes, S. C., Lama, T. M., Grutzner, F., Warren, W. C., Balakrishnan, C. N., Burt, D., George, J. M., Biegler, M. T., Iorns, D., Digby, A., Eason, D., Robertson, B., Edwards, T., Wilkinson, M., Turner, G., Meyer, A., Kautt, A. F., Franchini, P., Detrich, H. W., Svardal, H., Wagner, M., Naylor, G. J. P., Pippel, M., Malinsky, M., Mooney, M., Simbirsky, M., Hannigan, B. T., Pesout, T., Houck, M., Misuraca, A., Kingan, S. B., Hall, R., Kronenberg, Z., Sović, I., Dunn, C., Ning, Z., Hastie, A., Lee, J., Selvaraj, S., Green, R. E., Putnam, N. H., Gut, I., Ghurye, J., Garrison, E., Sims, Y., Collins, J., Pelan, S., Torrance, J., Tracey, A., Wood, J., Dagnew, R. E., Guan, D., London, S. E., Clayton, D. F., Mello, C. V., Friedrich, S. R., Lovell, P. V., Osipova, E., Al-Ajli, F. O., Secomandi, S., Kim, H., Theofanopoulou, C., Hiller, M., Zhou, Y., Harris, R. S., Makova, K. D., Medvedev, P., Hoffman, J., Masterson, P., Clark, K., Martin, F., Howe, K., Flicek, P., Walenz, B. P., Kwak, W., Clawson, H., Diekhans, M., Nassar, L., Paten, B., Kraus, R. H. S., Crawford, A. J., Gilbert, M. T. P., Zhang, G., Venkatesh, B., Murphy, R. W., Koepfli, K.-P., Shapiro, B., Johnson, W. E., Di Palma, F., Marques-Bonet, T., Teeling, E. C., Warnow, T., Graves, J. M., Ryder, O. A., Haussler, D., O’Brien, S. J., Korlach, J., Lewin, H. A., Howe, K., Myers, E. W., Durbin, R., Phillippy, A. M., and Jarvis, E. D. (2021). Towards complete and error-free genome assemblies of all vertebrate species. Nature, 592(7856), 737–746.

Wickham, H., Averick, M., Bryan, J., Chang, W., McGowan, L., François, R., Grolemund, G., Hayes, A., Henry, L., Hester, J., Kuhn, M., Pedersen, T., Miller, E., Bache, S., Müller, K., Ooms, J., Robinson, D., Seidel, D., Spinu, V., Takahashi, K., Vaughan, D., Wilke, C., Woo, K., and Yutani, H. (2019). Welcome to the tidyverse. Journal of Open Source Software, 4(43), 1686.

